# Stability of Alternative Bacteriorhodopsin Folds

**DOI:** 10.1101/2023.06.21.545983

**Authors:** Kristine A. Mackin, Douglas L. Theobald

**Affiliations:** Department of Biochemistry, Brandeis University, 415 South St., Waltham, MA

## Abstract

Bacteriorhodopsin is a light-activated proton pump found in archaea and some single-celled eukaryotes (Findlay and Pappin, 1986; Sharma et al., 2006; Spudich et al., 2000). This protein adopts the 7TM fold found used by all type I and type II rhodopsins. In previous work, we used bacteriorhodopsin from *Haloterrigena turkmenica* to demonstrate that substantially altered protein folds exhibit light-activated proton pumping (Kamo et al., 2006; Mackin et al., 2014). In this work, we further characterized these novel folds by assessing the stability of our mutants. We used SDS denaturation to calculate the change in unfolding free energy relative to the wild-type (Cao et al., 2012). We also determine the extinction coefficient for each mutant. These results demonstrate that even dramatic structural rearrangements do not critically destabilize the protein, although the extinction coefficient does vary independently of the stability and the proton-pumping activity. Interestingly, the position of the A helix in the protein sequence has the largest effect on the stability of the mutant; those mutants where A is not located at a terminus are destabilized compared to the wild-type.

## Introduction

Rhodopsins are integral membrane proteins that use the 7 transmembrane alpha helix fold, also found in G-protein couple receptors (Spudich et al., 2000). Two families of rhodopsins can be classified based on sequence similarity; the type I rhodopsins are found in prokaryotes and some single-celled eukaryotes (Sharma et al., 2009), and exhibit a range of activities including ion transport and sensory functions. Type II rhodopsins are primarily eumetazoan, and most function as photoreceptors (Heintzen, 2012). All rhodopsins of known structure adopt the seven transmembrane α-helix fold (7TM), despite of the insignificant sequence similarity between the two families. In addition, both families of rhodopsins have a conserved lysine position in the seventh (G) helix, that forms a covalent bond to the retinal cofactor responsible for the light-activated conformational change (Spudich et al., 2000).

This identical fold and other shared structural and functional features must be explained. Sequence information is insufficient to resolve the longstanding controversy between homology and convergence for these families; multiple analyses have arrived at opposing conclusions (Findlay and Pappin, 1986; Ihara et al., 1999; Larusso et al., 2008; Metzger et al., 1996; Oesterhelt and Stoeckenius, 1971; Pardo et al., 1992; Soppa, 1994; Spudich et al., 2000; Taylor…, 1993). A structural argument in favor of convergence hypothesizes that the shared fold is uniquely suited to perform the light-sensitive function, so independently evolving proteins would converge on the observed 7TM fold multiple times.

It was this structural prediction that we tested in the previous work, and by demonstrating that seven alternative folds are functional, we provide evidence in favor of divergence from an ancient common ancestor (Mackin et al., 2014). Complementary work on bovine rhodopsin demonstrates that the conserved lysine position, required for Schiff base formation with the retinal cofactor, can also be dramatically altered but still activate transducin in a light-dependent manner (Devine et al., 2013). This corresponds with our finding that a double mutant in bR retains proton pumping activity (Mackin et al., 2014).

Here, we further characterize the stability of our seven novel folds, and correct the reported rate of proton pumping using a measurement of the amount of photoactive pigment. In the previous study, the novel bR folds exhibited activity ranging from 40%-176% of the wild-type (Mackin et al., 2014). Correcting based on the ratio of photoactive pigment indicates that the rate of proton pumping varies even less than previously reported, further supporting the hypothesis that multiple folds can function as light-activated proton pumps. We found that all mutants are destabilized compared to the wild-type, and the magnitude of this change is largely attributable to the flexibility of the A helix. These results provide additional evidence that the 7TM bacteriorhodopsin structure is remarkably flexible and that multiple architectures are all functionally competent.

## Materials and Methods

### Bicelle Unfolding Experiments

Bacteriorhodopsin mutants were expressed and purified as described (Mackin et al., 2014). 1,2-Dimyristoyl-sn-glycerol-3-phosphocholine (DMPC) and 3((3-cholamidopropyl)- dimethylammonio)-2-hydroxy-1-propanesulphonate (CHAPSO) were purchased from Anatrace. Sodium dodecyl sulfate (SDS) was purchased from Fisher, and all-*trans* retinal (RET) from Sigma-Aldrich.

We used the bR_f_-to-bRO_u_ assay described Cao et al (Cao et al., 2012). A stock solution of 40 mM NaPi was prepared at pH 6.3. DMPC and CHAPSO were weighed to make a stock solution at 30 mM DMPC, 32 mM CHAPSO and NaPi and water were added to a final concentration of 20 mM buffer. This suspension was subjected to multiple freeze-heat-cool-sonicate-vortex cycles to form bicelles. SDS stock solution was prepared at 10% (w/v) in water. The bicelle solution was diluted with buffer containing 300 mM NaCl and 50 mM Tris at pH 6.5, adding protein to a final concentration of 100μg/mL (3.7μM), 10 mM NaPi pH 6.3, 15 mM DMPC, 16 mM CHAPSO, and 11.2 μM all-*trans* RET. Finally, SDS was added to a final concentration between 0 and 5%, with intervals determined by preliminary denaturation experiments to sufficiently characterize the transition zone.

Samples were equilibrated at room temperature for 5 days, then the absorbance of 500 μl of each sample was measured at 280 and 550 nm using the cuvette slot on a NanoDrop 1000-C. The percent SDS was transformed into χ_SDS_ as in (Cao et al., 2012), excluding water. This was graphed against the absorbance at 550 and fit using the equations below.

For unfolding experiments with added RET, the equation derived in Cao et al. 2012 was used:

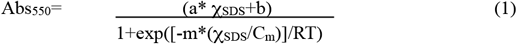

Parameters a, b, m, and C_m_ were fit using Kaleighdagraph 4.1.3. C_m_ is the concentration of SDS at the midpoint of the unfolding transition, and this value is used to compare the relative stability of the various novel fold mutants.

### Measurement of Extinction Coefficient

The hydroxylamine bleaching technique was used to determine the extinction coefficient of WT and mutant bR. We found that in the absence of added SDS the Schiff base was not accessible to the hydroxylamine, even after extended incubation of up to 7 days. Therefore, 3.0% SDS was added to bicelles of 20 mM DMPC, 32 mM Chapso bicelles with 150 mM NaCl, 20 mM NaP_i_, pH 6.3. 1M NH_2_OH, in significant excess over the protein concentration of 6 μM, was used to fully bleach the retinal. Control samples with neither NH_2_OH nor SDS were recorded and compared to the spectra observed after bleaching is complete. All samples were measured in the light-adapted state, as overnight dark adaptation did not have a significant effect on either the magnitude of the extinction coefficient or the λ_max_.

Previous work has established that the isomer of retinal present during the hydroxylamine attack is maintained (TREHAN et al., 1990), and this archaeal bR will isomerize between all-trans and 13-cis retinal. The extinction coefficient of these retinal oximes has been measured previously, and the average value of 52000 M^-1^ cm ^-1^ at 365 was used for our calculations (Imamoto et al., 1992; TREHAN et al., 1990). Since the protein is purified in the absence of added retinal, we assume that all retinal oxime present originated from cofactor bound to bR, and that therefore this concentration is equivalent to the amount of photoactive bR in the sample. This concentration is then used to calculate the extinction coefficient of each mutant, using the absorbance at 550 nm from the control spectra.

## Results

### Extinction Coefficient and Reconstitution Efficiency

The extinction coefficient of bacteriorhodopsin from *Haloterrigena turkmenica* (HtbR) has not previously been reported, so it was determined using the hydroxylamine reaction(Mukohata et al., 1991; Rehorek and Heyn, 1979).We relied on the reported extinction coefficient of retinal oxime at 365 nm from previous work, and used the average value of 52000 M^-1^cm^-1^ for the mixture of retinal oximes we expect after bleaching the light-adapted bR. This calculation gave an ε_550_ of 40000 (+/- 900) M^-1^cm^-1^, as compared to the reported ε_568_ 53000 M^-1^cm^-1^ of *Halobacterium salinarum* bR (HsbR) (Mukohata et al., 1981). The extinction coefficient for each mutant and the wild-type is reported in Table 1 as an average of three experiments. The standard error is also reported.

**Table 1:** χ SDS, and m from curve fits, ΔΔG_u_ values based on the midpoint of the wild-type unfolding transition. ΔΔG_u_ is reported is kcal mol^-1^. The value reported is the average of three trials with the standard error.

| **Mutant** | **$\chi$ SDS** | **m** | **$\Delta\Delta G$** |
| --- | --- | --- | --- |
| Wild Type | 0.59 (0.01) | -24 (4) | - |
| GBCDEFA | 0.56 (0.02) | -19 (1) | -0.4 (0.3) |
| BW | 0.553 (0.002) | -28 (2) | -1.1 (0.2) |
| CDEFGBA | 0.50 (0.01) | -26 (4) | -2.4 (0.2) |
| FW | 0.41 (0.01) | -15 (1) | -2.7 (0.3) |
| GFABCDE | 0.47 (0.02) | -28 (6) | -3.2 (0.3) |
| DW | 0.43 (0.02) | -21 (2) | -3.3 (0.3) |
| CDEFABG | 0.47 (0.01) | -28 (3) | -3.6 (1.0) |

The total concentration of protein in the sample was then calculated for each mutant based on the A_280_ (using the extinction coefficient calculated by the ExPASy server) (Arnold et al., 2012). The percentage of photoactive pigment was then determined by dividing the concentration calculated from the hydroxylamine reaction by the total amount of bR in the sample. This gives an estimate of the amount of functional protein present in each sample, and recapitulates the purity of the sample as calculated by a photometric analysis of an SDS-PAGE gel.

### Stability of Mutants

Using the equilibrium unfolding method described by Cao & Bowie (Cao et al., 2012), we recorded denaturation data for all of the mutants discussed in previous work. These data were fit with equation (1) to find the χ_SDS_ at the midpoint of the unfolding transition (C_m_). Preliminary results were used to approximate the C_m_, then subsequent experiments more fully characterized the unfolding transition zone (Figure 3). The m value, reported in Table 2, was used to calculate the change in unfolding free energy (ΔΔG_u_) of the mutants at the center of the wild-type unfolding transition (χ_SDS_=0.59).

**Table 2:** The extinction coefficient of mutants was calculated from the bleaching experiments, which were also used to calculate the approximate amount of photoactive protein in the sample.

*Stability of Mutants*
| **Mutant** | **$\epsilon_{550}$** | **% Photoactive Protein** |
| --- | --- | --- |
| Wild Type | 40000 (900) | 83 (4) |
| GBCDEFA | 43000 (2000) | 78 (3) |
| BW | 55000 (2000) | 53 (5) |
| CDEFGBA | 46000 (3000) | 35 (6) |
| FW | 51000 (1300) | 68 (2) |
| GFABCDE | 50000 (1500) | 67 (2) |
| DW | 58000 (3300) | 57 (1) |
| CDEFABG | 56000 (1500) | 40 (12) |

## Discussion

In this work, we were interested in further characterizing the previously described novel folds and determining what affect the fold reorganization has on the protein. The stability of bacteriorhodopsin has been examined extensively, due to the ease of observation based on the characteristic absorbance of the chromophore (Allen et al., 2001; Cao et al., 2012; Curnow et al., 2011; Faham et al., 2004). Previous work has examined the effect of changes in the inter-helical loops on the stability of the protein (Kim et al., 2001). However, the total reorganization of the primary sequence to generate a novel fold is, to our knowledge, unique, and we aim to understand how these changes affect the stability.

The change in unfolding free energy for the majority of the fold mutants examined here is negative, indicating that the stability is decreased relative to the wild-type. The GBCDEFA mutant is within error of the wild-type, and this any change in unfolding free energy is within one RT (0.592 kcal mol^-1^) indicating that this difference is likely insignificant. In contrast, the remainder of the mutants are definitively less stable than the wild-type.

The destabilized mutants appear to group loosely into two categories; those which are only slightly destabilized and those which are significantly destabilized, based on a cutoff of -2.5 kcal mol^-1^. Initially, we hypothesized that all of the mutants containing the artificial WALP helix would be similarly destabilized, especially considering that these mutants have the lowest rate of proton pumping. The relatively small change in unfolding free energy for the BW mutant therefore came as a surprise.

The stability of the mutants appears to be largely governed by one factor; whether helix A has one or two connections to the rest of the protein. In the less stable mutants, namely, CDEFABG, GFABCDE, DW, and FW, the A helix is connected on both ends of the peptide to the other helices. In contrast those that are not substantially destabilized only connect on one end of the A helix. Significantly, the BW mutant is the only WALP-containing mutant where the A helix is not bound on both ends. We propose that having two points of connection forces the A helix to make unfavorable contacts with the other helices, resulting in decreased stability. The change in unfolding free energy for the CDEFGBA mutant is within error of the numerical cutoff for the division between these two groups, in spite of the freedom of the A helix, and the increased instability of this mutant compared may be attributable to the alteration of the BC linkage. The least stable mutant, CDEFABG, suffers from both destabilizing features. The A helix is trapped between the F and B helices, placing it in the more destabilized category, and altered BC linkage adds an additional penalty.

Although the interhelical BC connection is not required for reconstitution of a functional pigment, changing or eliminating this native connection does have detrimental affects on the stability of the protein(Kahn and Engelman, 1992; Marti, 1998). The least stable mutant, CDEFABG, suffers from both destabilizing features. The A helix is trapped between the F and B helices, placing it in the more destabilized category, and altered BC linkage adds an additional penalty (Kataoka et al., 1992). Our observation recapitulates that of previous work by Booth and co-workers who found that altering the BC loop had the most significant impact on bacteriorhodopsin from H. salinarum (Kim et al., 2001).

We also measured the extinction coefficient of each mutant, and used this data to calculate the percent of photoactive pigment in the sample. The calculated value tracks closely with the purity of the sample as determined by analysis of commassie-stained SDS-PAGE gels, indicating that unfolded or otherwise inactive bacteriorhodopsin is not a large component of the sample. For future experiments, the extinction coefficient determined here can be used to calculate both the concentration and purity of bR samples.

Variability in extinction coefficient due to diverse changes in the protein sequence have been observed in previous work (Allen et al., 2001; Lu et al., 2001). Even dramatic changes made at the ends of the helices have been observed to maintain λmax identical to the wild-type, as with our previous work where the characteristic absorbance was largely unaltered from the WT, even with extreme changes in the protein fold (Figure 1) (Allen et al., 2001; Mackin et al., 2014). Experiments done with splitting the bacteriorhodopsin helices into two or more fragments, while recapitulating chromophore binding and proton-pumping activity, also altered the extinction coefficient (Kahn and Engelman, 1992; Kataoka et al., 1992; Marti, 1998). Changing the amino acid sequence of the loop also resulted in changes in the extinction coefficient while maintaining protein function and chromophore binding (Kim et al., 2001).

**Figure 1:**
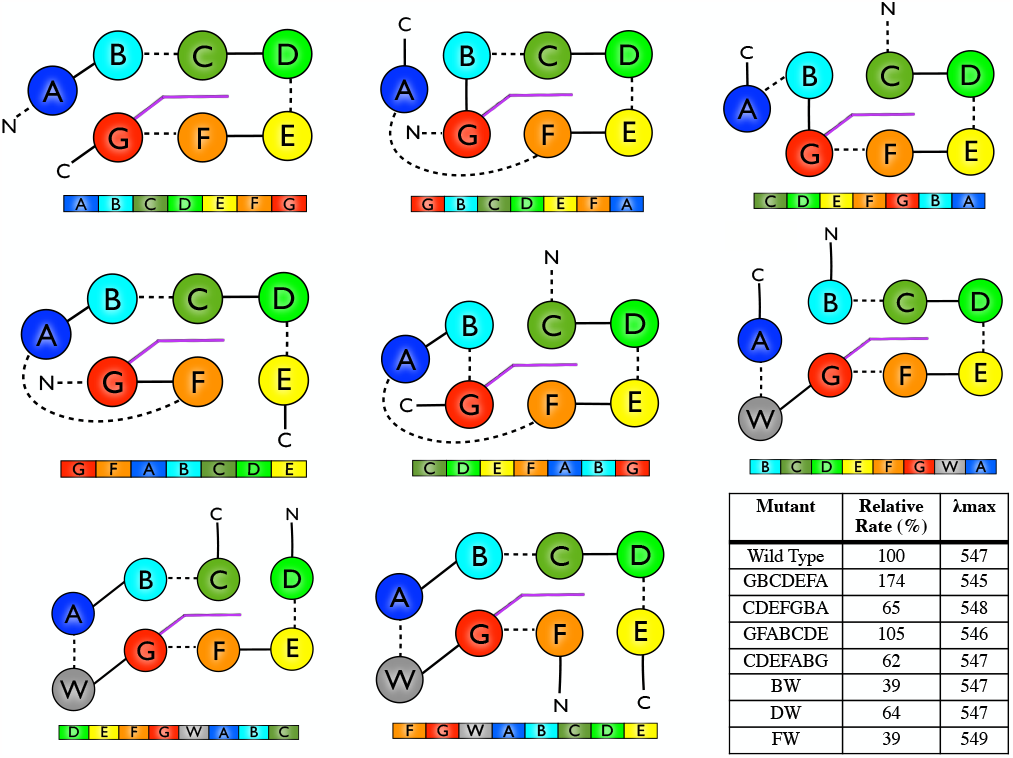
Bacteriorhodopsin permutation constructs, described and characterized in previous work (Mackin et al., 2014). All 7 novel folds support proton pumping activity, fold correctly and bind chromophore to display the characteristic absorbance. The rate relative to the wild-type is reported in the table.

**Figure 2:**
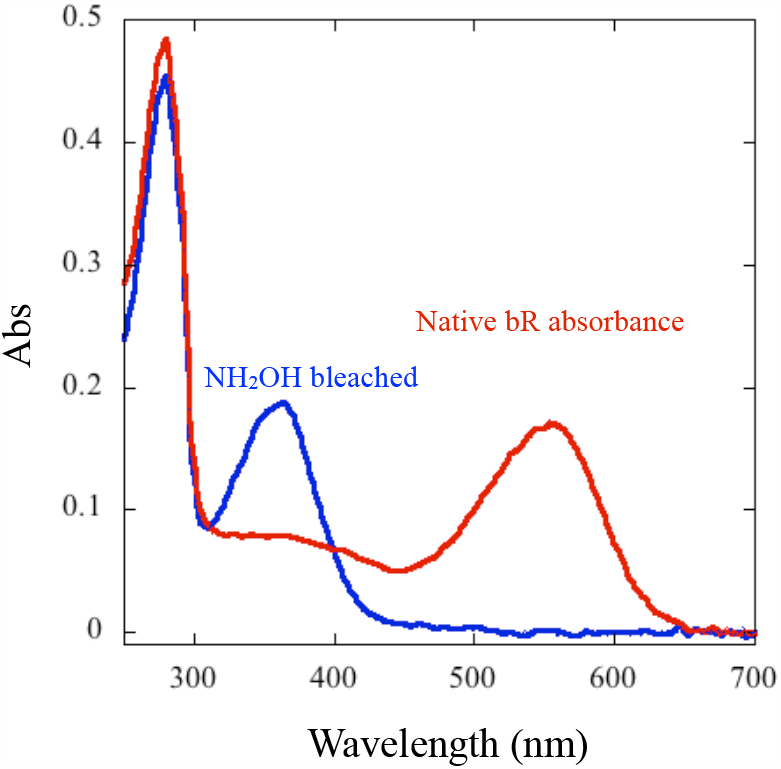
Spectra of the wild-type bacteriorhodopsin, showing the native absorbance (before bleaching) and the absorbance after bleaching with added NH_2_OH and SDS.

**Figure 3:**
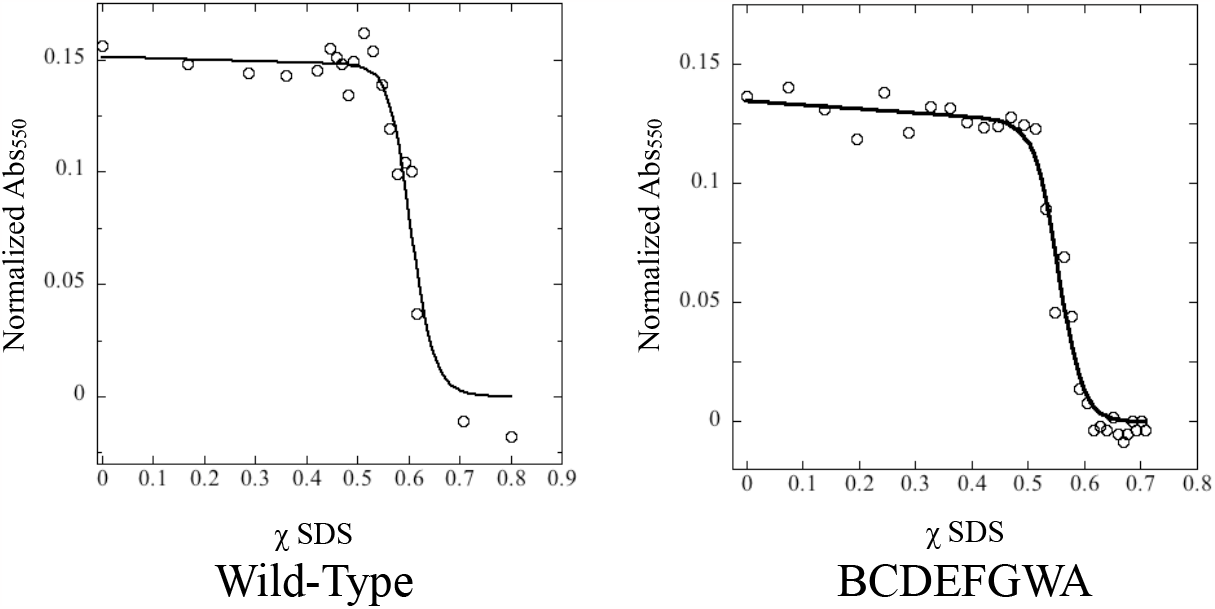
Representative examples of denaturation curves of the wild-type bR and BW mutant, showing the fit of equation (1) to the data.

**Figure 4:**
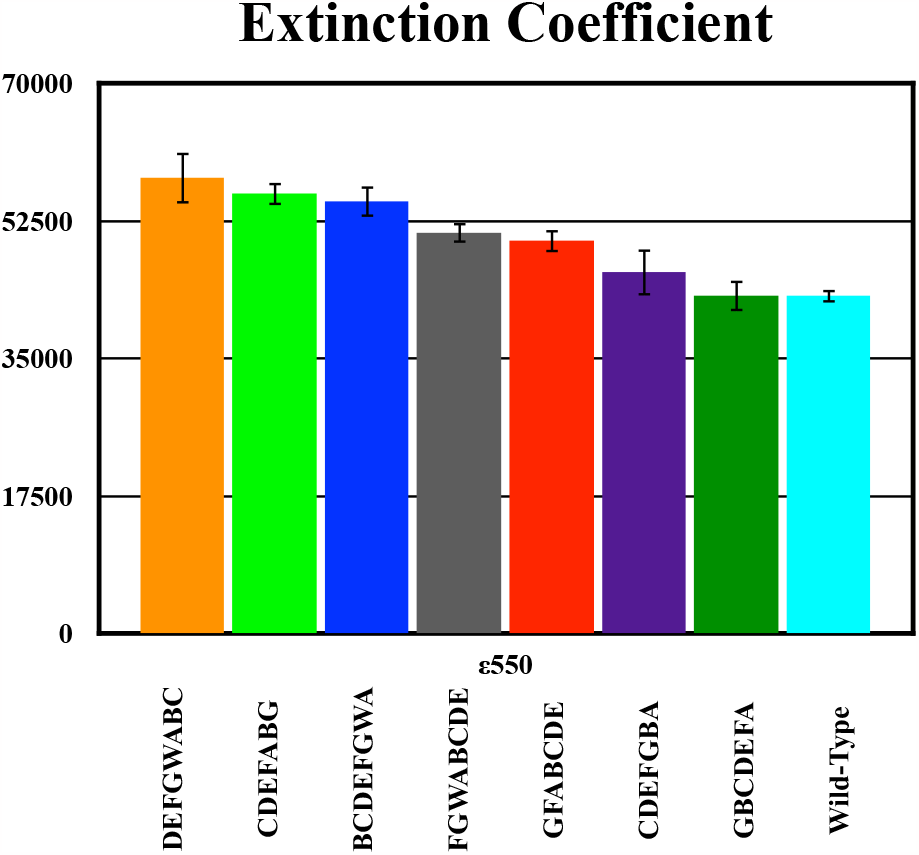
The extinction coefficient of each mutant; error bars represent the standard error from three experiments.

In previous work, the A280 was used to calculate the total amount of protein present in the sample. The rate of proton pumping observed in these experiments is thus a lower bound. Using the percent purity calculated here, which was consistent across at least two different preparations, we correct the previously reported rate to reflect the actual rate of protein pumping attributed to only photoactive bacteriorhodopsin. These rates are shown in Figure 5.

**Figure 5:**
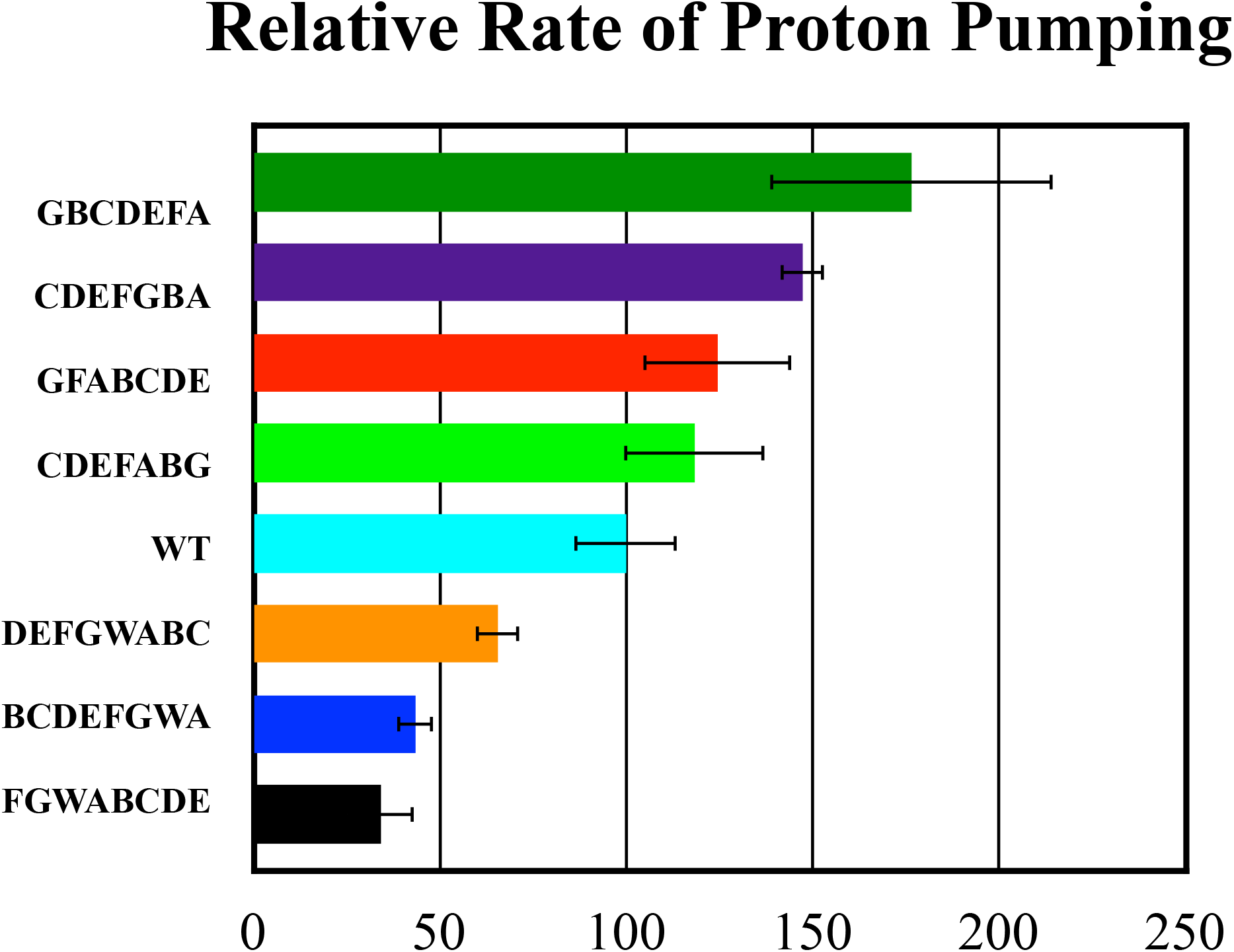
Relative rates of proton pumping, adjusted based on the percentage of photoactive protein present in the sample. Error bars are the standard deviation of multiple trials.

The results presented here do not demonstrate any significant relationship between the extinction coefficient and the rate of proton pumping. Interestingly the magnitude of this destabilization is neither correlated to the rate of proton pumping nor to the extinction coefficient. The only affect of the decreased stability we have observed is that increasingly destabilized mutants express at progressively lower concentrations in E. coli systems, although this relationship has not been quantified. Although the relative rate is slightly different based on this correction, these results do not substantially alter our conclusions.

